# The population dynamics of a bacterial pathogen after host re-infection affects the founding population size

**DOI:** 10.1101/061408

**Authors:** Gaofei Jiang, Rémi Peyraud, Philippe Remigi, Alice Guidot, Richard Berthomé, Wei Ding, Alexandre Jousset, Stéphane Genin, Nemo Peeters

## Abstract

In natura, many organisms face multiple infections by pathogens. The ability of a pathogen to reinfect an already-infected host affects the genetic makeup of the pathogen population at the end of the infectious cycle. Despite the likely prevalence of this situation, the population dynamics of pathogens during multiple infections over time is still poorly understood. Here we combined theoretical and empirical investigations of the founding population size, a critical driver of the evolution of pathogens, in a setting allowing for multiple and subsequent re-infections. Using the soil-borne bacterial pathogen Ralstonia solanacearum and tomato as its host, we first assessed the strength of the host infection bottleneck, and showed that both the host barrier and the immune system work additively to constrain the infection. Then, by increasing the temperature, we experimentally demonstrated that the increased pathogen proliferation within the host reduces the contribution of subsequent re-infection leading to a lower founding population size. Our study highlights the importance of within-host pathogen proliferation in determining founding population size – and thus bacterial genetic diversity during epidemics – for pathosystems where multiple re-infections occur. Under current global changes, our work notably predicts that an increased temperature provided this increase has a beneficial impact on pathogen growth, should decrease the founding population size and as a consequence potentially lower the diversity of the infecting and transmitted pathogen population.

**Significance Statement:** Founder population size is a major determinant of pathogen evolution, yet we still have limited insights into effective populations in natural settings. Most studies have considered infection as a single event, followed by pathogen growth in the host. But, in natura, organisms typically face multiple infections by several co-exisiting pathogen strains. Therefore, effective population size will depend on the timing and relative growth rate of the different infecting strains. In this work, we predict and experimentally show that both priority effects and within-host competition determines effective founding size, with an over-contribution of fast-growing and early infecting genotypes. This work sheds a new light on the ecological and evolutionary pressures affecting infection dynamics in realistic conditions.

## Introduction

Colonisation of new habitats by living organisms is often associated with a reduction of their population size. This variation of population size, also called bottleneck, can strongly shape their evolution in addition to natural selection^2^. The lower the size of the founding population in a new habitat, the more genetic drift will impact the evolutionary dynamics^45^. This is particularly important for pathogens that, after living and potentially growing outside the host, need to infect and proliferate within a new host habitat. They will encounter various layers of host physical and immune barriers or even competition from other microorganisms^2,6^. All these parameters may reduce the population size, making evolution stochastic, but may also provide a strong selective pressure that will drive pathogen evolution. Hence, assessing the factors influencing the founding population size of a given pathogen, provides key insights in the understanding of the genetic structure of their populations^7^ and should enable to better understand the complex life cycle of pathogens and provide means to better control them. However, to date most studies have extrapolated pathogen dynamics from single-infection models^1,3^. This is valuable but miss a key feature of epidemics, namely the high occurrence of multiple and recurrent infections.

Co-infections of a single host by multiple pathogen genotypes are common in nature and are an important driver of pathogen dynamics on humans^8^, animals^9
10^ as well as plant hosts^1^. The type of reported co-infections can be very different, ranging from infection by disctinctive pathogens; *e.g.* virus and bacteria^118^, to different strains (or genotypes) of the same pathogen^12,13
14^. Theoretical and empirical studies have investigated the pathogen population dynamics and the severity of the disease outcome upon co-infection^81512,14^. However, to our knowledge, little is known on the effect of multiple infections over time, of the same (re-infections^15^) or different pathogens (super-infections^16,17^ Gog *et al.*, 2015). This is in our opinion a major knowledge gap as multiple and successive infections situation may prevail in nature. For instance, animals face multiple pathogen exposures during recurrent social interactions, repeated feeding on contaminated food and repeated exposure to pathogen vectors^10,15^. For sessile organisms, like plants^12,14^, the environment can be a steady source of pathogen inoculum^18^. In such repeated infection settings, the assessment of the founding population size of pathogens to understand the global pathogen pathogen dynamics is missing. We can hypothesize that subsequent infections may lead to significant modification of the intra-host population composition following the arrival of new pathogen genotypes. On the other hand, the population dynamics of the first infecting individuals may modify the host environment and thus reduces or enhances the colonization success of the subsequent invaders^15^.

Our objective in this study is to determine the factors influencing the founding population size when a host is subjected to subsequent infections by the same pathogen. To this end, we combined mathematical modeling and experimental approaches. First, using a mathematical model, we explored how variation in the different key steps of the life cycle of a host-infecting bacterial pathogen in order to identify which parameters affect the size of the pathogen founding population when comparing single and multiple infection settings. We found that, besides bottleneck stringency, the post-bottlenecks population dynamics can also impact the founding population size when subsequent re-infection are considered.

Then, we empirically validate the model predictions by using tomato - *Ralstonia solanacearum* as model plant-pathogen system: The *Ralstonia* species complex is a highly diverse bacterial pathogen group that contains several devastating soil-borne pathogens with a broad host range^19^ and posing a serious threat to global crop production. The bacteria invade the host root xylem vessels and spread rapidly to aerial parts of the plant through the vascular system. In susceptible hosts, high population levels (up to 10^10^ colony-forming units per gram of fresh weight) are reached within less than a week, inducing vascular clogging causing wilting symptoms and plant death. Since these bacteria infect their hosts through roots and are known to be resilient in contaminated fields for decades^20^, we hypothesised that multiple re-infections could occur naturally in plant grown in contaminated soil. We empirically demonstrated that *Ralstonia sp.* strains inoculated onto the soil of their tomato host could indeed re-infect an already-infected host. Given that the pathogenic cycle is fast and reproducible upon simple soil-drenching inoculation, this plant-pathogen system is therefore well suited to experimentally investigate the role of the intra-host population dynamics on the founding population size when subsequent re-infections occur.

In this study, we determine, both through modeling and experimental tests, that the strongest contributors to the founding population size are the plant bottleneck (additive effect of both plant physical barriers and plant immunity) and, when subsequent re-infections are taken into account, within-host pathogen proliferation. By specifically manipulating within-host growth kinetics (by an increase of 2°C of the local environment), we show that the increase of the post-bottleneck proliferation of the pathogen is accompanied by a significant decrease of the founding population size. Our study demonstrates that under the re-infection model, growth within the host determines founding population size additionally to bottlenecks and that environmental changes modulate this dependency.

## Results

### Pathogen infection modes influence founding population size

We designed a mathematical model to theoretically explore the impacts of host-entry bottlenecks and within-host population dynamics on the founding population size during subsequent re-infections. This model reproduces the key steps of a microbial pathogen population infecting a host (Fig 1a, see Methods section). The first step corresponds to the transition from the pathogen environmental reservoir to the invasion of the host at the infection site, where the main bottlenecks occur. To allow for multiple re-infections, the establishment of the founding population is a continuous process concomitant with the subsequent steps: within-host proliferation, symptomatic phase and later transmission. This sequence of steps can vary according to the nature of the pathogen but describes particularly well a wide range of fast-growing and quorum-sensing dependent pathogens^21^, like it is the case for the plant pathogen *Ralstonia spp*.^22^.

**Figure 1.**
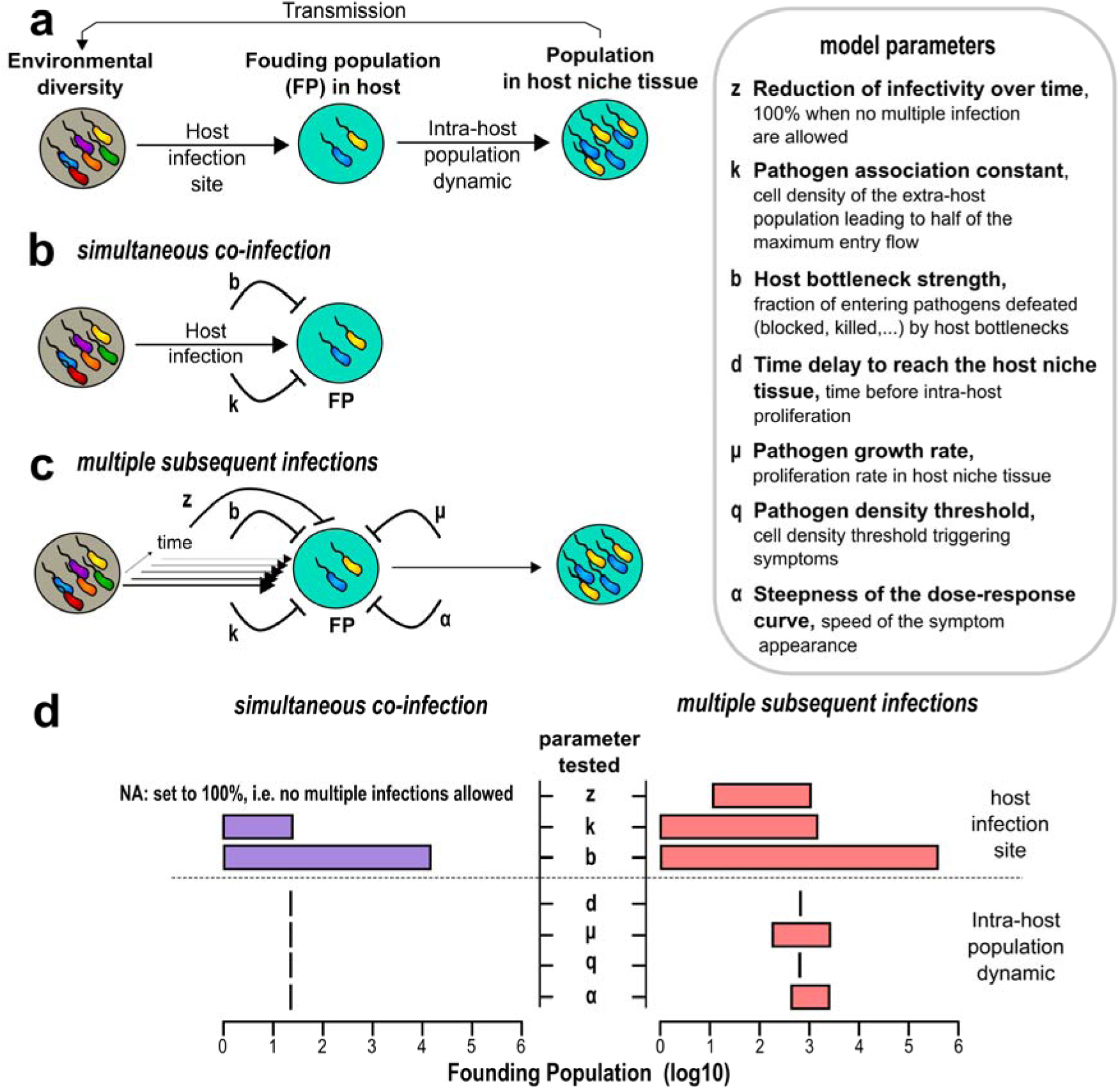
**Modeling of a bacterial pathogen infection process under variable infection modes.** a) Overview of two schematic steps modeled as regard to the founding population: entry within the host and the post-bottleneck infection dynamics. b) Influence of the main model parameters on the founding population in simultaneous multiple infections. Sign of repression-like indicates that the *FP* is reduced when the value of the parameter increase. c) Influence of the main model parameters on the founding population in subsequent multiple infections. d) Sensitivity analysis of the founding population size to model parameters in above two infection modes (b and c). The bar plot represents the maximum and the lower range of *FP* variation obtained by varying the parameter reported. Definitions of the main model parameters are listed in the box panel and Supplementary Note 1.

We then investigated which steps of the life cycle, from infection to transmission, have the most prevalent effect on founding population size. We considered either simultaneous co-infection or subsequent re-infections scenarios. Mathematical simulations with a z parameter for reduction of infectivity over time (Fig. 1c, 1d) revealed that upon subsequent infections, the bacterial population dynamic follow a very distinct variation compared to simultaneous infections. In a simultaneous co-infection, only the pathogen association constant (k) and the strength of the host bottleneck (b) influence the founding population size (Fig. 1b, 1d). The first parameter (k) represents the infection dose of the pathogen near the host infection sites required to penetrate the host. The second parameter (b) is indicative of the percentage of individuals blocked or killed by host factors driving the infection bottleneck, leading to a reduction of the pathogen entry flow. Allowing for multiple re-infections radically changes the contribution of the different model parameters to the founding population size. In the latter case, not only parameters governing the host infection (z, k and b), but also those controlling the kinetics of disease progression (µ and α), were found to affect the founding population size (Fig. 1c, 1d). Indeed, a higher growth rate (µ) of the pathogen inside the host niche tissue leads to an earlier transmission phase and thus reduces the number of pathogen cells entering from subsequent re-infections (Supplementary Fig. 1). The steepness of the dose response curve (α), is directly correlated to the growth rate (µ) as the quorum-sensing threshold triggering the expression disease symptom-inducing factors will be reached faster as the growth increases. A sensitivity analysis of the founding population size to each model parameter (see Supplementary Fig. 1 and Supplementary Note 1 for detailed results) identified the bottleneck (b) as the highest source of founding population size variability (Fig 1d) in either the simultaneous co-infection or the subsequent re-infection settings. Regarding the post-bottleneck infection dynamics, the growth rate µ of the pathogen within the host tissue also has a significant incidence on the founding population size but only in the subsequent re-infection setting (Fig. 1d).

### Infected tomato plants can be re-infected by Ralstonia pseudosolanacearum

The tomato-*Ralstonia sp.* pathosystem is ideal to test whether hosts can face multiple and subsequent re-infections by pathogens constitutively present in their surrounding microbiota. Indeed, this bacterium can survive in the soil whilst maintaining its virulence for years^23^. Furthermore, the soil-drenching inoculation procedure mimicks the presence of the bacteria in infested soils, allowing for a *quasi* natural infection. However, the actual possibility of multiple re-infections to occur in this pathosystem is unknown.

We performed non-simultaneous inoculations of tomato plants with two equally virulent strains of *R. pseudosolanacearum*, the wild type strain GMI1000 and a gentamicin-resistant derivative, GRS540. Importantly, both strains were shown to be of equal plant infectivity (Supplementary Fig. 2a, and^24^). Tomato plants were soil-drenched with strain GMI1000 and re-inoculated by soil drenching with strain GRS540 at 1, 8, 23 and 48 hours after the first inoculation (hours post-infection or hpi). We then tested for the co-occurrence of the GRS540 strain with the GMI1000 strain in the stem of each plant. With a re-inoculation at 23 hpi, 60% of the plants could be re-infected by GRS540 (Fig 2a). Re-infections can thus occur in plants grown in a contaminated soil. We found that the capacity of subsequent re-infection decreases over time at a rate, z, of 2.0 ± 0.1 %·h^-1^, with very few plants being re-infected at the late, 48 hpi, re-inoculation (Fig. 2a).

**Figure 2.**
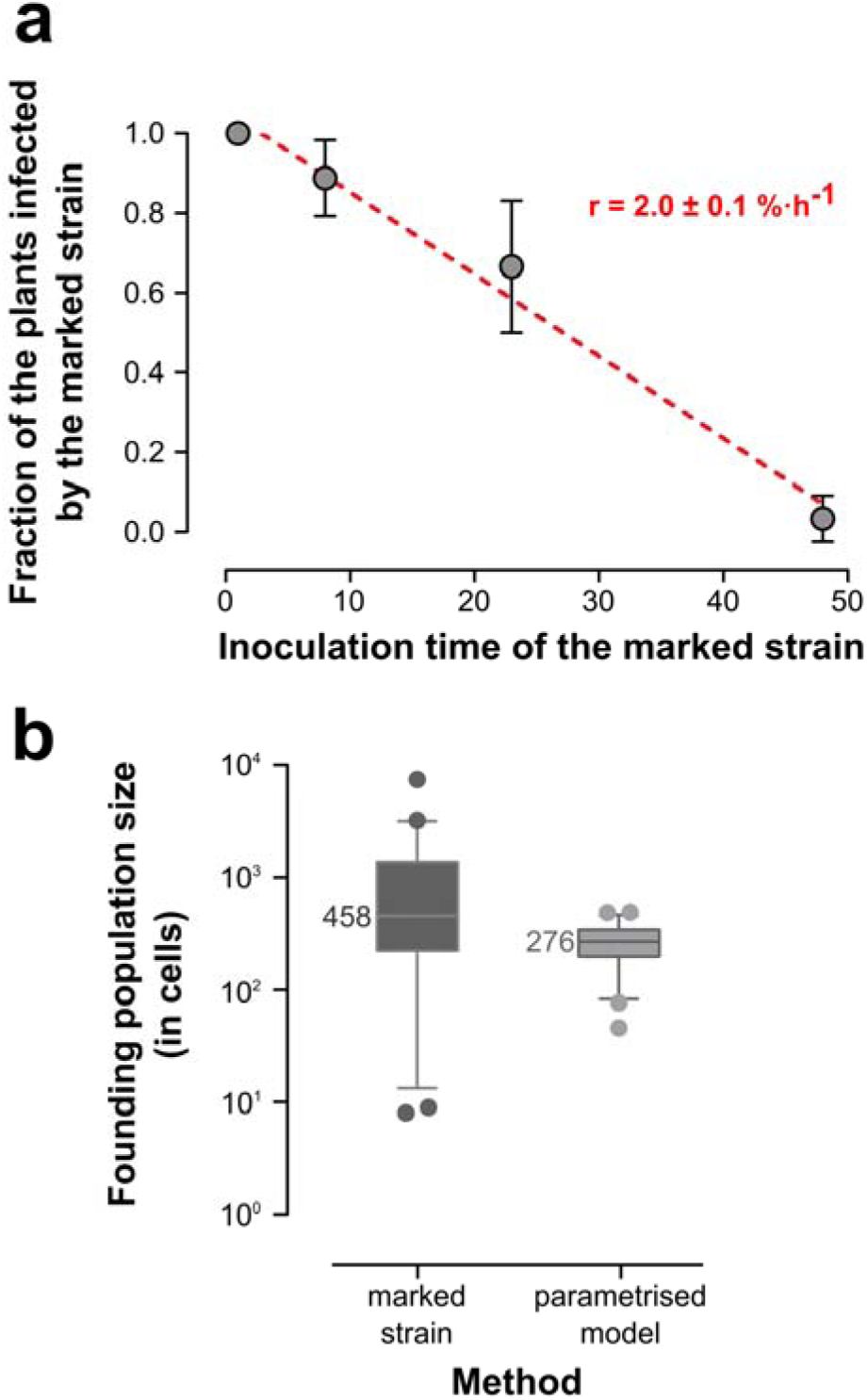
**Infection dynamics of *R. pseudosolanacearum* on tomato.** a) Infected tomato plants can be re-infected by R. pseudosolanacearum: Plants were first inoculated with the wild type GMI1000 strain and subsequently inoculated with the marked GRS540 strain (Supplementary Fig. 2a) at 1, 8, 23 and 48 hpi (hours post GMI1000 inoculation). Plants were havested at 5 dpi (days post-inoculation) to obtain the fraction of plant infected by GRS540. b) Measurement of the founding population size using the marked strain method and the model method (Methods section). Box plots represent the median and the 5-95 percentiles.

### The founding population size equals a few hundred cells

As tomato re-infection by *R. pseudosolanacearum* is possible (although within a short time frame of less than 48 hours), we thus favour the infection model where re-infection is allowed (with a z parameter of re-infection decrease over time). This model predicts contribution to the founding population size of both the bottleneck and the within-host population dynamics (Fig. 1d). We thus first sougth to evaluate the actual founding population (*FP*) size. We used two independent methods to quantify *FP* size, first an empirical method based on the co-infection with a mixture of two isogenic strains (‘marked strain method’), secondly a modeling method using the above established mathematical model (‘model method’).

For the empirical method, we used a statistical assessment of FP size based on the co-infection of a reference strain (GMI1000) mixed with minute fractions (ranging from 1% to less than 0.01%) of the isogenic marked strain GRS540. The probability of an infected plant to contain also the marked strain is dependent on the proportion of the marked strain in the original inoculum (*p*) and on the size of *FP* (see Methods and Supplementary Fig. 3). Using this method, we found a median *FP* size of 458 cells with a two times standard deviation of 2734 cells (Fig. 2b). The high variability of *FP* could reflect the true biological variability of the founding population; however, it may also arise from the stochastic sampling of the marked strain inherent to the low number of founding cells. We evaluated the contribution of a stochastic sampling by simulating a random sampling of the marked strain (see Methods). We identified that up to 18% of the variability observed (with a standard deviation of 507 on the 2734 observed) can be attributed to the sampling bias of the method indicating that most of the observed variability arises from true biological variability in founding population size (Supplementary Fig. 4).

Then, to quantify FP size using a modeling approach, we parametrised the model previously described (Fig. 1). This consisted in measuring and estimating the variability of the model parameters (Fig. 1; Methods section and Supplementary Note 2) during the infection process of a tomato plant by the reference strain GMI1000 (see Fig. 3). The lowest median infection dose (MID) indicates the infectivity of this pathogen on host surface, and can be assimilated to the pathogen association constant k (here equal to 5.9·10^6^ cells·ml^-1^ Fig. 3a, Supplementary Fig. 6, 7 and Note 2). Once inside the host, a time delay (d, 51 h ± 3h) is required for bacteria to reach their host niche; the plant xylem vessels for pathogenic *Ralstonia spp.* (Fig. 3b and Supplementary Fig. 8). Within that niche, bacteria will grow (rate µ of 0.233 ± 0.012 h^-1^, Fig.3c) to reach densities allowing quorum sensing-mediated production of virulence factors (here exopolysaccharides (T_n_)) with a given production flux (Supplementary Note 2). Symptom development follows a sigmoid curve with a steepness α of 0.612 and a scaling factor β of 2.31 (Fig. 3d and Supplementary Note 2). The first symptoms are trigged by EPS accumulation when the bacterial density reaches a threshold q of 6.0·10^7^ ± 0.1·10^7^ cfu·g_(FW)_-1__ (Fig. 3e and Supplementary Fig. 10 and 11). Following symptom development, *Ralstonia spp.* keeps growing, but with a reduced rate µ^-^ of 0.25 h^-1^ likely due to resource and space limitations (Supplementary Note 2).

**Figure 3.**
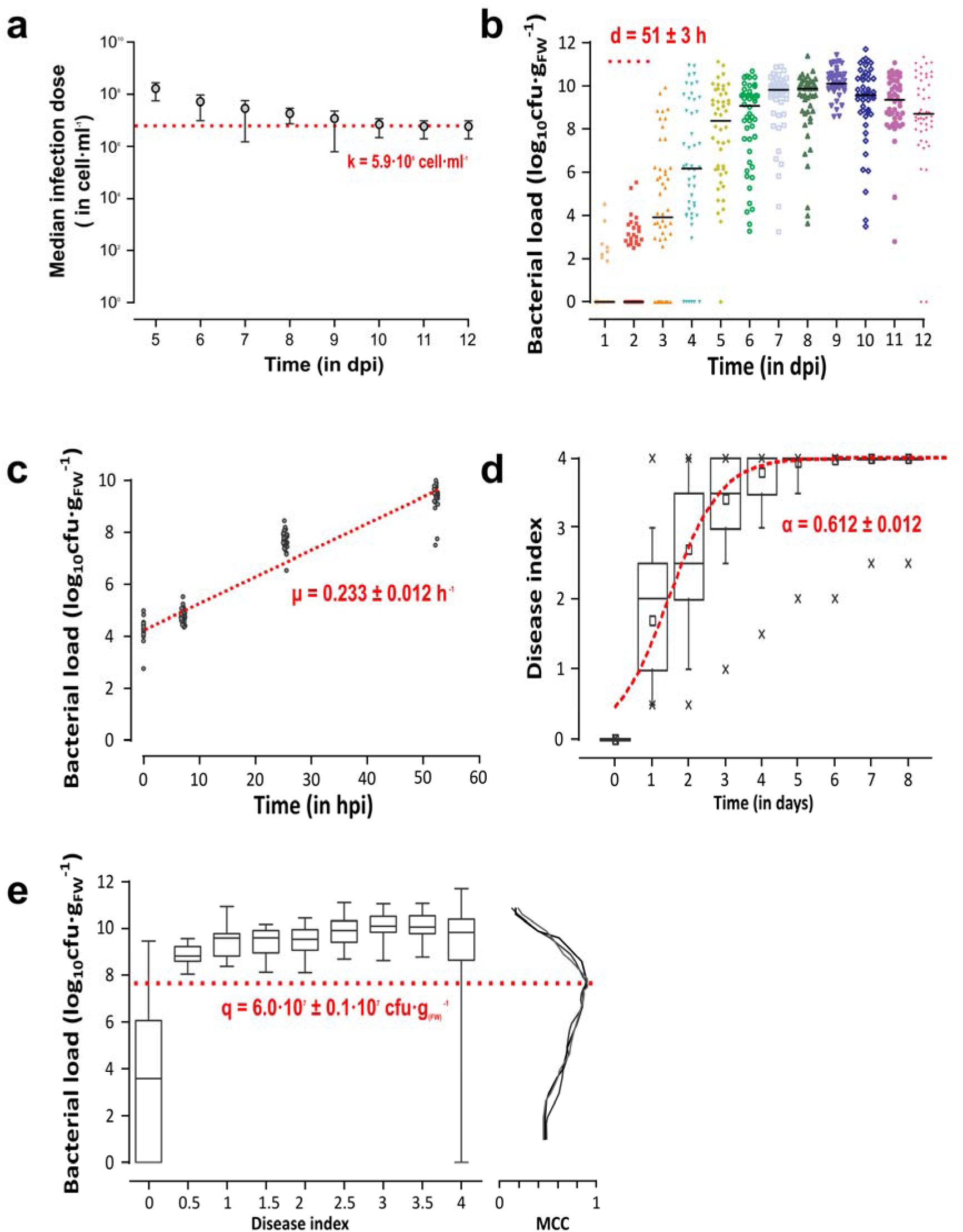
**Empirical assessment of the main parameters determining the outcome of the tomato – *R. pseudosolanacearum* interaction.** a) The median infection dose (MID) of *R. pseudosolanacearum* in tomato following soil-drenching inoculation. Daily phenotyping was performed (Supplementary Fig. S3) and the Reed-Muench method^36^ was adopted to determine the MID at each dpi. The red dotted line represents an asymptotic value at MID = 5.9·10^6^ cells which is a good proxy for the association constant, K, between the plant surface and the pathogen. b) Colonisation dynamics of *R. pseudosolanacearum* in xylem. Bar represents the median of bacterial load in each group. The red dotted line represents the delay time, d at 51 hours, to reach the niche value. c) *In planta* growth of *R. pseudosolanacearum* after stem injection inoculation. Thirty-two individual plants from 8 biological replicates were analysed. The red dotted line represents the fitted exponential growth rate, µ, with a value of 0.233 ± 0.012 h^-1^; R^2^ = 0.937± 0.024 (mean ± sd). d) Time response curve of symptom onset (wilting) synchronized to the first day of the symptom onset for each plant. The red dotted line represents the best fit of a wilting dose response. With the x marks representing the 95% confidence interval e) Relation between bacterial wilt and disease index. The data from Fig. 3b and Supplementary Fig. 10 was used for this analysis. Box plots represent the median and 5-95 percentiles in each DI group. The red dotted line represents the quorum sensing threshold, q, inducing the production of virulence factors responsible to the host symptoms. A value of 6.10^7^ ± 0.97.10_7_ cfu·g_(FW)^-1^_ was found to be the best predictor of the wilting onset based on Matthews correlation coefficient (MCC) (see Supplementary Fig. 11).

Then, we used the predictive capacity of the parametrised model to determine the founding population size. The calculated value of the founding population size at 7 dpi is of 276 ± 105 cells, in the experimental conditions used to define the parameters of the model. This value was in the range of the observed value using the marked strain methods (*FP*_WT_ = 458 cells; see Fig. 2b), under the same experimental conditions as used for the parametrisation.

### Plant physical barrier and immunity are major bottlenecks

Our previous theoretical model identified the host bottleneck as the major driver of the founding population size. In order to assess experimentally the contribution of the bottleneck stringency in controlling the *FP* size, we manipulated two main plant defense mechanisms: the root physical barriers and the immune system. First, we quantified *FP* size using a root wounding inoculation procedure to estimate the contribution of root physical barriers. At 4 dpi, all root-wounded plants had symptoms, indicating an accelerated infection and colonization. We used the marked strain method to estimate the founding population size in wounding condition (*FP*_w_). The size of *FP*_w_ dramatically increased to a median of 24 215 cells (Fig. 4), significantly higher than the founding population size for non-wounded plants (*FP* = 458 cells) as previously defined (Mann Whitney test, P-value < 0.0001).

**Figure 4.**
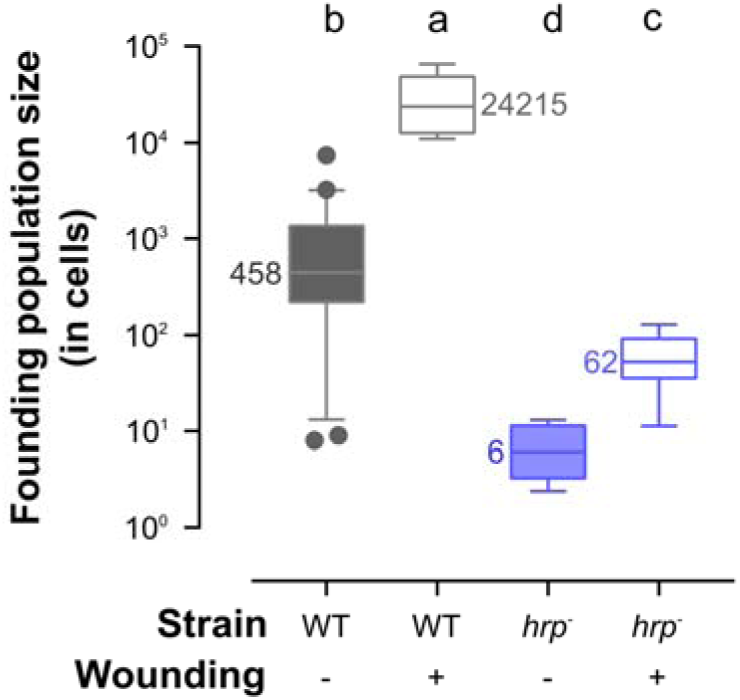
**Contribution of the plant physical barriers and the plant immunity to the infection bottleneck.** Founding population size measured empirically for wild type GMI1000 of R. pseudosolanacearum (grey) and hrp mutant GMI1694 (blue). GMI1000 and GMI1694 strains were harvested at 7 dpi and 10 dpi following root inoculation respectively. GMI1000 wounding (WT-wounding) and GMI1694 wounding (hrp-wounding), both in white boxes, were harvested at 4 dpi and 7 dpi by wounding root inoculation. Different letters above the box plot indicate a Mann Whitney test, P-value < 0.0001.

In order to evaluate the bottleneck effect due to the plant immune system clearance, we inoculated tomato plants by soil drenching with a *hrp* mutant (GMI1694) and its isogenic gentamycin-marked strain (GRS743; see Supplementary Fig. 2b). GMI1694 is an *hrcV* mutant devoid of Type 3 secretion system^25^, and thus unable to suppress plant immunity triggered upon infection^26^. The corresponding founding population size, *FP*_hrp_, was experimentally estimated at a median of 6 cells (and mean of 7 ± 4 cells, Fig. 4), significantly lower than the wild-type strain (*FP*_WT_ = 458 cells) as previously defined (Mann Whitney test, P-value = 0.0014).

The predicted and the measured *FP* size data obtained previously (Fig. 4) enabled us to assess the relative strength of the two bottleneck factors: the plant physical barrier and the immune system. First, we fitted the high values (*FP*_w_ = 24 215 cells) found in the root-wounding experiment and the earliest symptoms observed (4 days) and found a high founder entry flow of 1065 cell·h^-1^. Hence, we estimated the strength of the bottleneck, b, due to the root physical barrier to be of 0.021 by comparing the founder entry flow of 22 cell·h^-1^ without wounding and 1065 cell·h^-1^ with wounding (see Supplementary Note 2 for details). This indicates that only 2.1% of the soil bacterial cells can pass through the root barrier, corresponding to a reduction of 97.9% of the soil bacterial cells at the root entry. Using the mathematical model (see Supplementary Note 2), the founder entry flow for the *hrp* mutant was estimated to be of around 1 cell·h^-1^ without wounding and around 3 cells·h^-1^ with wounding. The strength of the bottleneck due to the immune system clearance was estimated to be around 0.002, equivalent to a clearance of 99.8% of the soil bacterial cells at the root entry.

Both the plant physical barriers and the immune system are involved in the bottleneck occurring at the entry site of the pathogen into the host. This could result in a potential bias in the quantification of the bottleneck triggered by each mechanism. To evaluate this potential overlapping effect, we quantified the founding population size of the *hrp* strain infecting wounded-root plants. Here, the *FP*_w-hrp_ size increased to a mean of 62 ± 40 cells and a median of 53 cells (Fig. 4). A *hrp* mutant entry flow of 3 cells·h^-1^ was fitted in this experiment (see Supplementary Note 2).

As the *hrp* mutant displays no specific phenotype *ex planta* (*in vitro* growth, motility), the defect in the Type 3 secretion system only affecting host-dependent phenotypes, we can hypothesise that the maximum entry flow (V_max_) in wounded roots of the *hrp* mutant should be similar to the wild type strain entry flow (1065 cells·h^-1^). As the successful founder flow corresponds to the survivors to the immune system response, we show that the strength of the immune system clearance is of 99.7% in this experiment. Thus, only 0.3% of the bacteria can escape the plant immune system. This is similar to the non-wounded experiment (immune system clearance of 99.8%) suggesting that both plant physical barrier and immune system factors works additively in the plant infection bottleneck excluding synergistic interactions.

### Within-host growth drives the founding population size

Theoretical assessment of the parameters driving the founding population size revealed that, under subsequent re-infections, not only the host bottleneck but also the post-bottleneck population dynamic defines the pathogen founding population size. As the growth of the plant pathogenic *Ralstonia sp.* is known to be strongly influenced by temperature, we tested the impact of a 2 °C increase (30 °C *versus* 28 °C, see Supplementary Note 1) on *FP* size. Such a temperature shift increases *in vitro* growth rate by 16% (t-test P-value= 0.0177), from 0.161 h^-1^ ± 0.030 at 28°C to 0.186 h^-1^ ± 0.020 at 30 °C (Fig 5a). The *in planta* growth rate at 30°C also increased, by 43% (t-test P-value= 0.004); from 0.233 h^-1^ ± 0.012 at 28 °C to 0.333 h^-1^ ± 0.014 at 30 °C (Fig 5a). Using these values as a new growth rate (µ) parameters in the mathematical model, we predicted a reduction in the founding population size from 276 ± 105 cells at 28 °C to 173 ± 68 cells at 30°C (Fig. 5b), corresponding to a reduction of 37.3%. Experimentally measuring the size of the founding population at 30°C using the marked strain method yielded a founding population size of 114 ± 76 cells at 30°C (Fig 5b). This value, which is close to the predicted one (173 ± 68), corresponds to a drastic reduction of 75%, compared of the measured founding population size at 28 °C (Fig. 5b). Therefore, post bottleneck population dynamics can significantly affect founding population size when subsequent re-infections occur.

**Figure 5.**
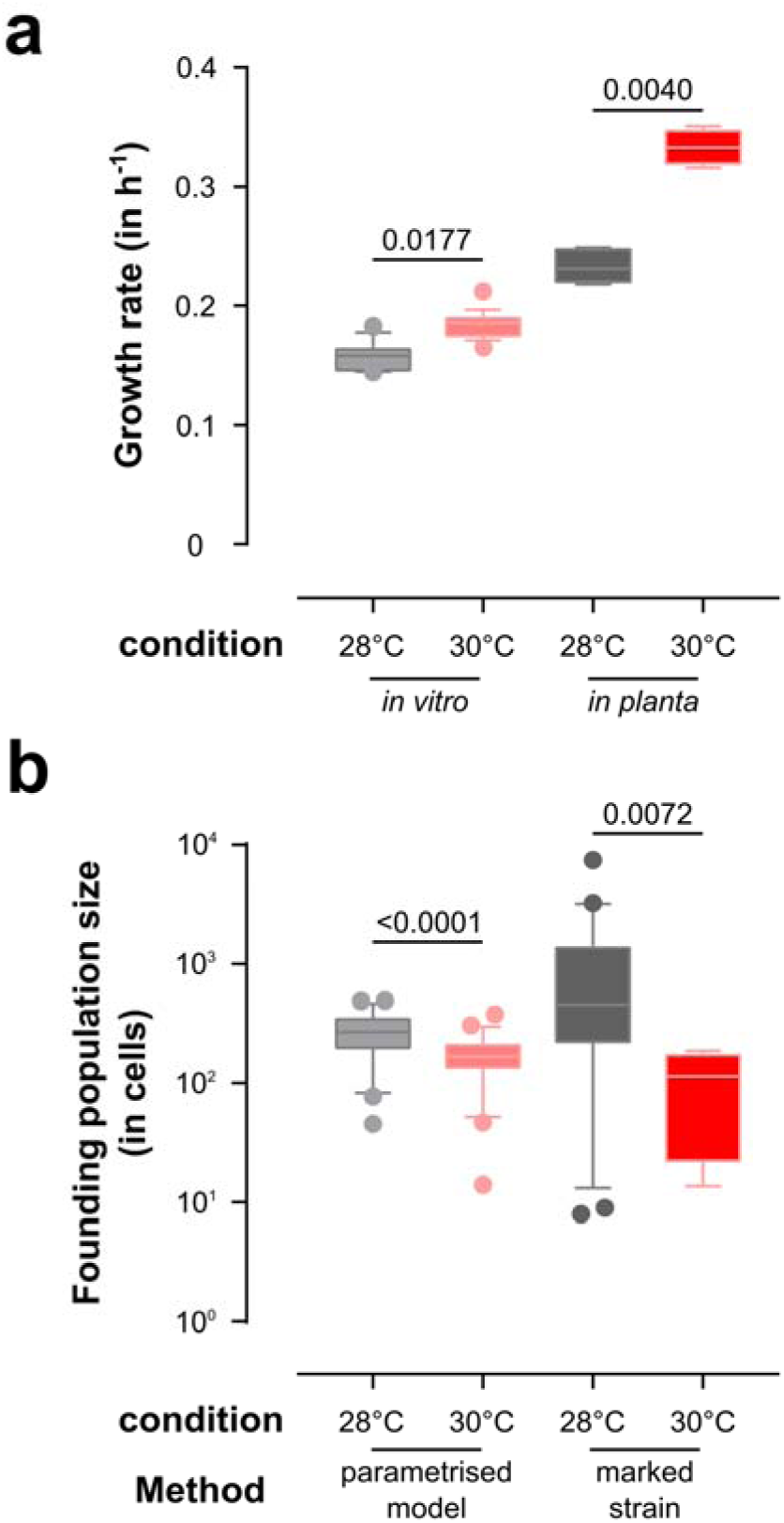
**Post-bottleneck bacterial growth drives the founding population size.** a) *In vitro* and *in planta* growth rate of *R. pseudosolanacearum* at 28 °C and 30 °C (Supplementary Fig. 9). b) The founding population size predicted by the parametrised model and measured by the marked strain method (Methods section) at 28°C and 30°C. numbers above the box plots indicate P-values from a Mann Whitney test.

## Discussion

To understand epidemics from an evolutionary perspective of the pathogen it is essential to identify the key determinants of the pathogen population size. At the transmission phase of an infection cycle, when the pathogen population returns to the local environment, a specific enrichment of within-host amplified genotypes will occur, with important epidemiological and evolutionary consequences^3^. A current challenge in the study of microbial pathogens is to understand how, and to which extent, within-host processes will affect the desiminated pathogen population^27–29^. One of these parameters is the size of the pathogen founding population, an essential factor shaping pathogens’ population at dissemination. Indeed, the reduction in genetic diversity of microbial populations infecting a new host, also call founder effect, may lead to genetic drift^30^.

Here we show that, in a plant-pathogen system where multiple subsequent re-infections can occur, the size of founding populations depends not only on the stringency of the infection bottleneck, but also on the rate of pathogen proliferation within the host, highlighting the importance of the sequence of infection. Our work demonstrates and measures that there is a slow-down/arrest of the infection process (including re-infections) when the pathogen population reaches a given threshold, at which symptoms develop and dissemination begins. Indeed, a higher proliferation rate leads to an earlier pathogen transmission. A large number of parasites may follow a similar infection strategy, including other hemibiotrophic plant pathogens^21^. Results from our mathematical model indicate that the effect of microbial proliferation during the infection on the founding population size should be a general characteristic of such pathogens. We believe that this work could be basis for re-evaluating other host-pathogen interactions (for which subsequent re-infection occur^1^) both in laboratory and field conditions. In this work, considering the experimental system used (R. pseudosolanacearum - tomato), we evaluate that the founding population size would have been underestimated by 20 times if re-infections had not been considered.

Indeed, most previous studies aiming at determining the size of pathogen founding population ignored the complexity of natural infection of a host in which a significant proportion can be considered as multiple re-infections^31,32^. In our experimental system, we found that there is a time window (up to 48h after the first infection) for potential subsequent re-infections of an already-infected host. The observation that the re-infection efficacy decreases rapidly over time could be due to plant immune responses^33^. This could also be due to a decline of rhizosphere nutrient availability, local modification in the root architecture or modification in the plant chemotaxis signals attracting the pathogen. As expected, the size of the pathogen founding population increased significantly after mechanically wounding the plant roots, as this lifts the natural barriers of the plant. We also showed that this size dramatically decreased when plants were infected with a hrp mutant able to colonize the plant without producing any symptoms^33^. This is explained by the incapacity of this bacterial mutant to suppress plant immunity. Our combined modeling and experimental approach allowed us to quantify the effect of these two bottleneck factors: taken separately they drastically limit the infecting population by 97.9% (plant physical barriers) and 99.8% (plant immunity). We were able to show that these two factors are additive in the bottleneck effect. The different locations where both bottlenecks occur, i.e. epidermis for physical barrier and inner cortex for immune system, could explain the additive effect observed instead of a synergistic effect. At a smaller scale, there could be specific bottlenecks for the bacteria to cross several distinct cell layers within the root organ^34^, like it was shown for the progression of the anthrax bacteria in different murine tissues^35^.

After the strength of the infection bottlenecks, the second most important contributor to the founding population size is the immediate within host proliferation, to be taken into account as subsequent re-infections are possible. Indeed, modeling of the infection cycle indicates that a higher within-plant growth rate (µ) of the pathogen has a negative impact on the size of the founding population. Using the 2°C increase of temperature as a mean to enhance bacterial growth, we show that the increased within-plant growth is accompanied by a significant reduction in the size of the founding population. The decrease of the founding population could also be due to an increased bottleneck associated with increased temperature, but this is unlikely as increased temperatures generally have a detrimental effect on plant immunity^36,37^ and would thus lessen this specific bottleneck. Furthermore, this small temperature shift should not affect the plant root integrity, ruling out a significant contribution of the plant physical barriers to this lowering of the founding population. The α parameter (speed of symptom appearance), is directly correlated with growth (µ), as the time to reach symptoms will be reduced with increased growth, and as such is also highlighted as an important contributor to the founding population size (Fig 1d).

The Ralstonia sp. complex has a wide species-to-continent geographic distribution^38^. Some diversity analysis of single isolates from different infected plants show that different genotypes from plant pathogenic Ralstonia species^38^ can be present over a narrow geographical distribution^39,40^. At the scale of a single field, high genotypic diversity could be identified among the Ralstonia sp. strains isolated from different plants^41^.

Our study opens new perspectives for studying contribution of the root resident or soil microbiome to the infection bottleneck as well as the impact of global environmental changes on the R. solanacearum spp.-plant interaction. As re-infections by R. solanacearum spp are possible, we demonstrated that the final genetic makeup is a combination of bottlenecks and bacterial population dynamics within the host. More detailed studies on the local soil diversity in plant pathogenic Ralstonia sp. are needed to evaluate the complexity of the potential virulent population in the rhizosphere soil. This would allow us to evaluate effective population size^5,42^ and thus the potential founder effect of the infection bottleneck and within host population dynamics under subsequent re-infection. Our work notably predicts that with increased temperature under current global changes, the founding population size should decrease and, as a consequence, this will potentially lower the diversity of the infecting and transmitted pathogen population. Overall, our study sheds light on the importance to consider re-infection to understand evolution and epidemiology of microbial pathogens. By demonstrating how in such situation a slight increase of local temperature can affect the founding population size of the pathogen.

## Methods

### Mathematical modelling of host infection

We designed a mathematical model to describe the population size of a pathogen (P_c_) and its founder number (F_c_) within the host. The model is composed of 3 compartments (c), the external compartment (e), the infection site (s), and the niche tissue (n). The volume dimension of the model was set in ml. P_c_ and F_c_ can be in the magnitude of few cells in tissues, thus we used entier value for these variables. The model mimicks the bacterial population dynamics in 5 steps.

Step 1) We modeled the flow of pathogenic cells entering at the infection site and facing host bottleneck from a contaminated environment. Pathogen cells in the environment (P_e_) reach the infection site with a probability depending on their environmental concentration, this is modeled by attributing an association constant of the pathogens to the host surface, (k). Secondly, infection founders at the infection site (F_s_) are cells which overcame constitutive and active defenses and then will disseminate into the host. Thus, the amount of infection founder depends on the maximum flow of pathogen cells entering without resistance, *i.e.* without bottleneck, (V_fmax_) reduced by the size of bottlenecks, constitutive and active defense, (b). Finally, the reduction of the flow of subsequent infections (due to acquired immunity or resources depletion) is captured by the decay, z, of a percentage of V_fmax_ by hour. Thus, the equation 1 gives the amount of infection founders crossing the infection site over time (t):

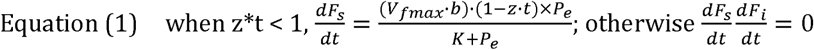

Step 2) Pathogen migrates from the infection site toward the niche tissue, where they will proliferate, may require some delay (d, in hour). So, the variation of founders in the host niche tissue (F_n_) is obtained from the following relationship:

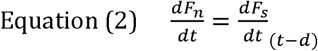

Step 3) The founding population proliferates in the niche tissue. The size of the population in the niche, P_n_, is dependent on the flow of the founding cells entering into this tissue F_n_, the growth rate of the pathogen in the niche tissue, µ in h^-1^, and the reduction of the proliferation, µ^-^, due to the collapse of the host tissues when the host symptom S (in arbitrary disease index, ranging from 0 to S_max_) increase during the course of infection.

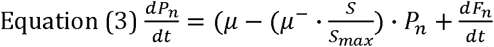

Step 4) Transmission of pathogens relies on the production of virulence factors. This production is often dependent on the cell density, when the pathogen population exceeds a quorum sensing system threshold^33^, q. Then the concentration of a toxin in the host, T_n_, in mmol·ml^-1^, is calculated from the flux of toxin production, V_t_.

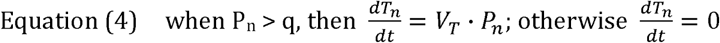

This equation may also mimick a population size exceeding the load capacity of the host when the transmission is not triggered directly by virulence factor production.

Step 5) The symptoms, S in disease index units (DI), is calculated by a dose response equation describing the steepness of the curve (parameter α and β) and the maximum symptom reached, S_max_, which depends on the toxin concentration in the host, T_n_.

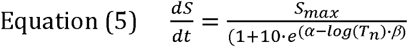

The equations are solved numerically with a time step of 1 hour. This time step was tested to be small enough to avoid numerical drift of the solution. The transmission time was defined as the time when the symptom reaches S > 85% S_max_. When no symptom level above 85% S_max_ is reached after 30 days, this time point is taken for the evaluation of the founding population size.

### Sensitivity analysis of the model parameters

To assess the influence of the model parameters on the founding population size we performed a sensitivity analysis in two conditions: 1) simultaneous co-infection where the parameter z is set at 1; 2) multiple subsequent infections where the z parameter is set < 1. The influence of each model parameter was tested by varying their values (the range is displayed in Supplementary figure 1) while others stay as default: k = 1·10^7^; V_max_ = 5·10^4^; b = 0.0002; d = 50; q = 1·10^8^; V_T_ = 1·10^-9^; µ = 0.2; µ^-^ = 0.2; α = 2; β = 0.5; S_max_ = 4; the concentration of the pathogen population outside the host, P_e_, was set to 1·10^7^.

### Bacterial strains, plant material and culture conditions

The *R. pseudosolanacearum* strains used in this work were: GMI1000, the wild-type strain; GRS540, a derivative strain from GMI1000 with a gentamicin cassette insertion^43^; GMI1694, a Type 3 secretion system mutant (*hrcV* mutant, *hrp*) with spectinomycin cassette insertion^25^; GRS743, a derivative strain from GMI1694 with the integration of the gentamicin cassette of GRS540 (this study). The gentamicin cassette does not alter the pathogen fitness, as was previously shown by stem injection^43^ and Supplementary Fig. 2. *R. pseudosolanacearum* strains were routinely grown at 28 °C in complete BG medium^44^. Antibiotics were used at the following final concentrations: 10 μg·L^-1^ gentamicin and 40 μg·L^-1^ spectinomycin. *Solanum lycopersicum* var. Super Marmande was cultivated in greenhouse. Square trays (30 30 4.5 cm) were used to grow 16 tomato seedlings and kept in the greenhouse under the following conditions: 75% humidity, 12 h light 24 °C, and 12 h darkness 22 °C. Plants were inoculated after 4 weeks of growth in dedicated quarantine facilities.

*In vitro* growth rate assays were performed in 200 μl liquid culture using 96 well plate under agitation at 600 rpm with FLUOstar Omega microplates reader (BMG Labtech). Bacterial cells were inoculated at a starting optical density 600 nm of 0.05.

### Experimental determination of the founding population size

A ‘marked strain method’ was developed to quantitatively estimate the infection founding population size of *R. pseudosolanacearum* in tomato, or *FP*_i_. The method is based on the measurement of the proportion of plants infected with a gentamycine marked strain when the host was inoculated with a mixture of the wt strain plus various small (0.1% to 0.001%) proportions of a marked strain. The starting frequency of the marked strain in the inoculum, *p*, was measured by serial plating. The frequency of the wt strain is 1 - *p*. *R* is referred to as the event that at least one cell of the resistant bacteria individual is present in the host plant. *P*(*R*) is the probability of the event *R. NR* is considered as the event that only the wt (Non-Resistant) strain is present. *P*(*NR*) is the probability of the event *NR*. Thus, *P*(*R*) + *P*(*NR*) = 1. Based on the above: *P*(*NR*) = (1 - *p*)^*FP^P^*^; *P*(*R*)=1 - (1 - *p*)*^FP^* and *FP* = log(1 - *P*(*R*)) / log(1 - *p*). Supplementary Fig. 9 highlights the sigmoid feature of *FP* = *f*(*P*) for each given *p. FP* or ‘infection founding population size’ can thus be estimated from this expression through investigating *P*(*R*) experimentally as follows: the resistant, marked strains are GRS540 or GRS743 (Supplementary Fig. 2), and their respective non-marked isogenic strains are GMI1000 (wt) and GMI1694 (*hrp*). The sampling time was at 7 dpi for root inoculation of GMI1000 and GRS540, 4 dpi for wounding inoculation of GMI1000 and GRS540, 10 dpi for inoculation of GMI1694 and GRS743 and 7 dpi for wounding inoculation of GMI1000 and GRS540. Stem samples were collected as described before and appearance of the two strains was determined by plating on BG agar with and without antibiotics. The two most extreme output value of *FP*, *i.e. FP* = 0 and *FP* = infinity, respectively obtained when *P*(*R*) = 0/32 or *P*(*R*) = 32/32 were excluded from the analysis as artificially skew the estimation of *FP*, indeed these limit values are the two asymptotes of the sigmoid curve of *FP* = f(P).

### Stochastic sampling simulation

Using the ‘model method’ we evaluated the contribution of stochastic sampling of the marked strain on the variability of founding population observed bymeasurement using the marked strain methodology. We simulated a random sampling of the cells based on the same parameters as the experimental setup. These parameters are the number of plants tested per replicate, 32, the concentration of marked strain in the inoculum 0.1% to 0.001% (*p*), and a founder population size of 458. A random number between 0 and 1 was generated for each 458 cell sampled, and the cell was assigned to be a marked strain if the number is below *p* in the inoculum. Then, simulated founding population size was calculated as the marked strain method above. Simulated values are shown in the Supplementary Fig. 4.

### Median infection dose

Wild-type *R. pseudosolanacearum* GMI1000 was used to determine the median infection dose (MID) in tomato. Groups of 16 tomato plants in one tray were soil drenching inoculated with 5·10^8^, 5·10^7^, 5·10^6^, 5·10^5^ or 5·10^4^ cfu·ml^-1^ in 500 ml of bacterial suspension. For stem injection, the inoculums were diluted to 5·10^7^, 5·10^6^, 5·10^5^, 5·10^4^ and 5·10^3^ cfu·ml^-1^ of *R. pseudosolanacearum* suspension and 10 µl bacterial suspension was directly injected into the stem with a microsyringe (Hamilton, Reno, NV, U.S.A.) at the cotyledon level. Symptom appearance was scored daily for each plant. Disease index (DI) was used to describe the observed wilting: 0 for no wilting; 1 for 25% of leaves wilted; 2 for 50%; 3 for 75% and 4 for complete wilting. Symptom onset (DI > 0) was defined as an infection success of *R. pseudosolanacearum* in tomato. The median infection dose (MID) of this pathogen in tomato was used at the 50% endpoint method of Reed and Muench formula^45^. Each treatment had two technical repeats and three biological repeats with 96 plants in total.

### Bacterial colonization kinetics and *in planta* growth rate

The soil-drenching inoculation was performed using GMI1000 at 5·10^7^ cfu·ml^-1^. Symptoms were recorded before harvest as described above. Sixteen plants were sampled at each dpi and this experiment was repeated three times with 48 samples in total. For the stem injection procedure, each plant was injected with 10 µl bacterial solution containing 5·10^6^ cfu·ml^-1^ of GMI1000. Sampling was conducted straight away after injection (0 hpi), 7 hpi, 24 hpi and 52 hpi. Each sampling had 4 plants and this experiment was repeated 8 times (32 samples in total). The 1 cm stem from the cotyledons upwards was harvested, weighed, and homogenized by mechanical disruption in a grinder (Retsch France, Verder, 95610 Eragny sur Oise, France). Bacteria were then enumerated by plating serial dilutions on BG agar plates, using an easyspiral (Interscience, Saint Nom la Breteche, France).

### Prediction of the bacterial load threshold inducing wilting

The following relation was used to predict wilting onset W (W = 1) or no witling (W = 0) depending on the bacterial load b, in cfu·ml^-1^, *in planta*. W = 1 if b > t then W = 0 with t, the threshold above which the wilting onset occurs. Two-class contingence matrices with experimental and simulated data were generated for changing values of t. The Matthews correlation coefficient (MCC)^46^ was used to compare predicting capacities to experiment values, also see Supplementary Fig. 10 and Note 2. The MCC can range from 0 (random assignment) to 1 (perfect prediction).

## Acknowledgements

GJ was supported by China Scholarship Council (No. [2013] 3009) and TECHNO II Erasmus Mundus Programme (2015-2016). RP was supported by EMBO (Long-Term Fellowship ALTF 1627-2011), Marie Curie Actions (EMBOCOFUND2010, GA-2010-267146), and European Research Council (ERC-StG336808 project VariWhim). This work was supported by the “Laboratoire d’Excellence (LABEX)” TULIP (ANR-10-LABX-41). The funding bodies played no role in study design, data collection and analysis, decision to publish, or preparation of the manuscript.

## Authors Contribution

GJ, RP, PR and NP designed the research; GJ, RP performed experiments; GJ, RP, AG, SG and NP analysed the data; RB contributed data; RP, GJ, PR, AG, AJ, SG, WD, RB and NP wrote the manuscript. All authors have read and approved the manuscript for publication. GJ and RP contributed equally to the work.

